# Using DeepLabCut-Live to probe state dependent neural circuits of behavior with closed-loop optogenetic stimulation

**DOI:** 10.1101/2024.07.28.605489

**Authors:** Melissa Gonzalez, Mark A. Gradwell, Joshua K Thackray, Komal R. Patel, Kanaksha K. Temkar, Victoria E. Abraira

## Abstract

**Background:** Closed-loop behavior paradigms enable us to dissect the state-dependent neural circuits underlying behavior in real-time. However, studying context-dependent locomotor perturbations has been challenging due to limitations in molecular tools and techniques for real-time manipulation of spinal cord circuits.

**New Method:** We developed a novel closed-loop optogenetic stimulation paradigm that utilizes DeepLabCut-Live pose estimation to manipulate primary sensory afferent activity at specific phases of the locomotor cycle in mice. A compact DeepLabCut model was trained to track hindlimb kinematics in real-time and integrated into the Bonsai visual programming framework. This allowed an LED to be triggered to photo-stimulate sensory neurons expressing channelrhodopsin at user-defined pose-based criteria, such as during the stance or swing phase.

**Results:** Optogenetic activation of nociceptive TRPV1^+^ sensory neurons during treadmill locomotion reliably evoked paw withdrawal responses. Photoactivation during stance generated a brief withdrawal, while stimulation during swing elicited a prolonged response likely engaging stumbling corrective reflexes.

Comparison with Existing Methods: This new method allows for high spatiotemporal precision in manipulating spinal circuits based on the phase of the locomotor cycle. Unlike previous approaches, this closed-loop system can control for the state-dependent nature of sensorimotor responses during locomotion.

**Conclusions:** Integrating DeepLabCut-Live with optogenetics provides a powerful new approach to dissect the context-dependent role of sensory feedback and spinal interneurons in modulating locomotion. This technique opens new avenues for uncovering the neural substrates of state-dependent behaviors and has broad applicability for studies of real-time closed-loop manipulation based on pose estimation.

**Manuscript:** *Highlights:*

1. Closed-loop system probes state-dependent behaviors at pose-modulated instances
2. Bonsai integrates DeepLabCut models for real-time pose estimation during locomotion
3. Phase-dependent TRPV1^+^ sensory afferent photostimulation elicits context-specific withdrawal responses

## 1. Introduction

State dependent behaviors in animals are influenced by internal or external factors that alter activity in a context dependent manner. Locomotion is a context dependent motor behavior enabling animals to navigate their environment by means such as walking, flying, or swimming. The dynamic nature of locomotion is facilitated by sensory inputs and spinal interneurons that modulate motor neuron activity to influence the temporal and alternating coordination of flexor and extensor muscles (Engberg & Lundberg, 1969; Grillner & El Manira, 2020; Harnie et al., 2022; Rossignol, 1996). To achieve coordinated and adaptable movements, these motor outputs must be responsive to incoming sensory inputs, such as unexpected external obstacles that may perturb locomotion. The stumbling corrective reflex, a flexor response in the swing phase and an extensor response in the stance phase, exemplifies this sensorimotor integration (Forssberg, 1979; Forssberg et al., 1977; Mayer & Akay, 2018; Wand et al., 1980). Studying the context dependent role of locomotor perturbations has been challenging due to molecular and technical limitations. This study aims to bridge this gap by providing a framework to couple nearly instantaneous pose tracking with pose modulated optogenetic stimulation at different phases of the step cycle.

Mice are excellent models for studying the neural mechanisms of locomotion due to the availability of genetic tools that allow for targeted manipulation of specific cell populations (Sjulson et al., 2016). Combining these genetic approaches with optogenetics enables researchers to control neuronal activity with high spatiotemporal precision (Boyden et al., 2005; Deisseroth, 2011). Optogenetic photostimulation occurs in real-time when light-sensitive proteins, expressed in genetically defined neuronal populations, are activated or inhibited by light (Yizhar et al., 2011). This technique offers the ability to target specific neurons at precise times, without inducing long-term compensatory mechanisms or neural rewiring that may occur with other techniques such as neuronal ablation or chemogenetics (Roth, 2016). Optogenetic manipulation of spinal cord neurons has been challenging due to the limited penetration of light through tissue. However, recent advances in optogenetic tools, such as the development of red-shifted opsins (Chuong et al., 2014; Klapoetke et al., 2014) and highly sensitive opsins (Mardinly et al., 2018), have enabled the manipulation of deeper populations of spinal interneurons. By integrating optogenetics with genetically encoded tools in mice, researchers are starting to dissect the role of specific neural circuits in locomotion with unprecedented specificity and temporal resolution (Kiehn, 2016; Kiehn et al., 2010).

Advances in open-source machine learning techniques for recording and quantifying animal behavior have paralleled improvements in the precision of cell targeting for refined neuronal manipulation. Supervised machine learning-based pose estimation tools, such as DeepLabCut (DLC) and SLEAP, require minimal human labeling to train a network to detect and track animal postures (Mathis et al., 2018; Pereira et al., 2022). DLC is particularly advantageous due to its ability to reliably capture user-defined features such as individual hindlimb joints, using high-performance feature detection and sophisticated deep learning models to analyze hindlimb kinematics (Mathis et al., 2018; Nath et al., 2019). The exceptional tracking performance of DLC becomes even more valuable when integrated into event-based frameworks like Bonsai, which allow for the processing of multiple data streams in real-time (Lopes et al., 2015). By incorporating DLC pose tracking, Bonsai enables the creation of a closed-loop feedback system capable of triggering an LED pulse at specific pose-modulated instances, opening up new possibilities for studying the neural mechanisms underlying behavior (Kane et al., 2020; Lopes et al., 2015).

The confluence of genetic targeting, spatiotemporal manipulation, and machine learning has created an unprecedented opportunity for closed-loop feedback in neuroscience research. By leveraging these powerful tools, we can now investigate the causal relationships between neural activity and behavior with unparalleled precision and specificity. In this study, we demonstrate the feasibility of this approach by instrumenting DeepLabCut-Live (Kane et al., 2020), a real-time pose estimation system, and optogenetics to manipulate a specific population of primary sensory afferents during locomotion in mice. By using DeepLabCut-Live to track the animal’s hindlimb kinematics and trigger optogenetic stimulation at specific phases of the locomotor cycle, we can probe the context-dependent role of these sensory afferents in modulating motor output and uncover the neural mechanisms underlying adaptive behaviors, such as the stumbling corrective reflex. This closed-loop system opens up new avenues for investigating the neural circuits underlying state-dependent behaviors and highlights the immense potential of integrating cutting-edge techniques from genetics, optogenetics, and machine learning in neuroscience research.

## 2. Materials and Methods

### 2.1 Animal subjects and ethical approval

TRPV1^Cre^;Advillin^FlpO^;R26^LSL-FSF-ChR2^ ^(Ai80)^ mice were used to target nociceptive primary afferent fibers. Mouse lines: TRPV1^Cre^ (JAX#017769)(Cavanaugh et al., 2011), R26^LSL-FSF-ChR2^ ^(Ai80)^ (JAX#025109)(Daigle et al., 2018), and Advillin^FlpO^ (Zimmerman et al., 2019).

Animal housing, surgery, and behavioral experiments conformed to Rutgers University’s Institutional Animal Care and Use Committee (IACUC: protocol #:201702589). All mice used in experiments were housed in a regular light cycle room (lights on from 08:00 to 20:00) with food and water available ad libitum.

### 2.2 Joint labeling for video tracking

Animals were anesthetized with 1.5% - 2% isoflurane to remove fur over the right hindlimb and right-side of the abdomen. Using White Oil-Based Paint Marker (Sharpie), dots were placed directly onto the skin over five anatomical landmarks: iliac crest (IC), hip, ankle, metatarsophalangeal joint (MTP), and the tip of the second toe. DLC is a markerless posture tracking system, but explicitly marking the hindlimb joints improves the ease of manual labeling and model training speeds. Due to dermal slippage, the knee joint is not marked. Instead, the lengths of the femur and tibia bones were measured to triangulate the position of the knee post-hoc given the 2D coordinates of the hip and ankle joints.

### 2.3 Treadmill locomotion training and video recordings

Two days prior to experimentation, mice were trained once each day to locomote on the treadmill by gradually increasing the belt speed from 5 - 20 cm/s. On experimentation day, animals were habituated to the behavior room for at least thirty minutes prior to placement in the Digigait™ motorized treadmill (Mouse Specifics, Inc., Boston, MA). Once accustomed to treadmill locomotion at average walking speeds of 20 cm/s, at least 5 step cycles were captured perpendicular to the treadmill using a Promon U1000 monochrome high-speed camera (510113-00-0000, AOS Technologies AG, Switzerland). The AOS Technologies Imaging Studio software suite was utilized on a high performance Windows machine to control camera capture with 864 x 796 pixel resolution at 415 frames per second (FPS). Infrared lights were placed on either side of the treadmill behind the camera to illuminate the scene and reduce glare (**Table 1**).

**Table 1.**
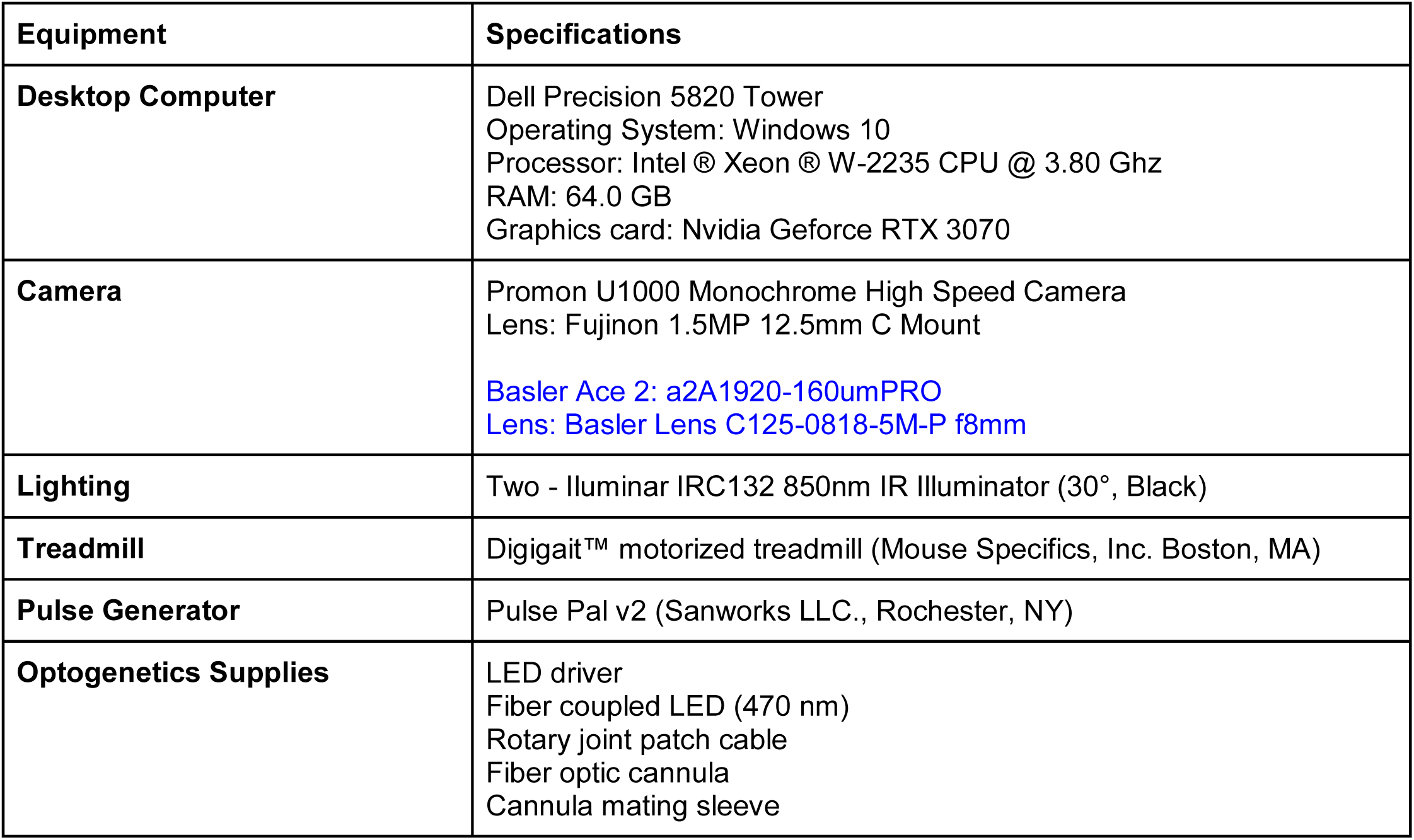
Hardware specifications for offline and real-time pose estimation as outlined in blue.

### 2.4 Fiber optic probe implantation, optogenetic stimulation, and c-Fos validation

Optic probe implantation was performed at 8-12 weeks of age. The optogenetic probe surgery was performed as previously described (Smith et al., 2019). In brief, animals were anesthetized with isoflurane (5% initial, 1.5-2% maintenance) and shaved over the thoracolumbar region. While positioned in a stereotaxic frame, a longitudinal incision (∼3 cm) was made over the T10-L1 vertebrae. The paraspinal musculature was removed to expose the spinal vertebra and the space between T12 and T13 was cleared to expose the spinal cord. Surgical staples were attached to T12 and T13 to provide a fixation point for the optic probe. A 400 nm core, 1 mm length fiber optic probe (Thorlabs), was then positioned over the exposed spinal cord, lateral to the midline, and secured to the surgical staples using layered dental cement (Ivoclar) and super glue (Krazy glue). The surgical site was closed with surgical staples, and the animals were allowed to recover for two weeks prior to behavior experimentation (**Figure 1A)**.

**Figure 1.**
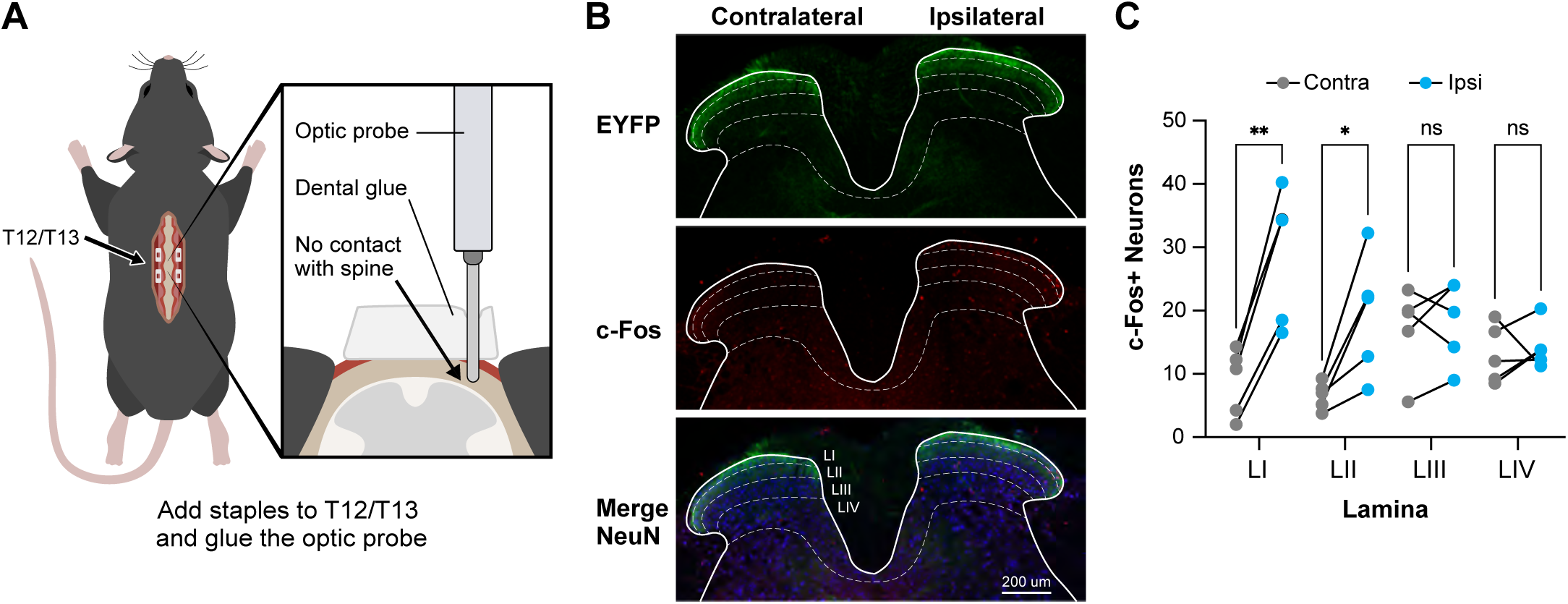
Probe implantation and validation of TRPV1^+^ afferent fiber activation. **(A)** Schematic of optic probe implant. **(B)** Transverse lumbar spinal cord section of TRPV1^Cre^;Advillin^FlpO^;^R26LSL-FSF-ChR2(Ai80)^ mouse following ipsilateral photostimulation. Top: EYFP (green), Middle: c-Fos (red), Bottom: Merge with NeuN (blue). Scale bar: 200 µm. **(C)** Quantification of c-Fos^+^ neurons by lamina contralateral and ipsilateral to photostimulation. Two-way ANOVA with the Geisser-Greenhouse correction and Sidak’s multiple comparison test. LI **p = 0.0016; LII *p = 0.0231, LIII ns p = 0.6637, LIV ns p = 0.6422. n = 5 mice.

To deliver LED stimulation, a Python compatible pulse generator, Pulse Pal (Sanworks) was connected to an LED driver (Thorlabs), which was in turn connected to a fiber-coupled LED (Thorlabs) (**Table 1**). The Pulse Pal was programmed to stimulate at 2.2 mW for 10 ms upon receiving a trigger signal from the host computer.

Optogenetic activation of TRPV1^+^ afferent terminals was assessed by delivering spinal photostimulation to anesthetized TRPV1^Cre^;Advillin^FlpO^;R26^LSL-FSF-ChR2^ ^(Ai80)^ mice and then processing spinal cord for c-Fos. Mice were anesthetized for 1 hr prior to photostimulation. Unilateral photostimulation (2.2 mW, 10 ms pulses, 10 Hz for 30 min) was delivered to the spinal cord by positioning an optic fiber probe (400 nm core, 1 mm fiber length, ThorLabs) above the spinal cord surface. Following photostimulation mice remained under anesthesia for a further 1 hr before transcardial perfusion with heparinized-saline followed by 4% paraformaldehyde in PBS. Tissue was dissected and post-fixed in 4% paraformaldehyde at 4°C for 2 hrs. Transverse sections (50 µm thick) were collected using a vibrating microtome (Leica VT1000S) and processed for immunohistochemistry as previously described (Hughes et al., 2012). Sections were incubated in a cocktail of primary antibodies: chicken anti GFP (1:1000, Aves), rabbit anti c-Fos (1:1000, Synaptic Systems), and mouse anti NeuN (1:2000, Millipore). Primary antibody labeling was detected using species-specific secondary antibodies (1:500). Sections were incubated in primary antibodies for 72 hrs and in secondary antibodies for 12-18 hrs at 4°C. All antibodies were made up in a 0.1M phosphate buffer with 0.3M NaCl and 0.3% Triton X-100.

### 2.5 Training DLC model for offline and online tracking

For offline tracking of hindlimb body parts, a DeepLabCut model was trained using a ResNet-50 backbone. A training dataset was prepared by adding 9 videos with an average of 2210 overall frames and extracting an average of 125 frames per video to manually label the five anatomical landmarks of interest. The dataset was shuffled and split 95:5 for training and testing, respectively. The network was trained for 300,000 iterations using the training subset, and then evaluated on the held-out testing subset. Using a confidence threshold of 0.9, we observed an average test error of 2.86 pixels and average train error of 2.67 pixels, compared to human provided annotations.

Following this procedure, an additional model was trained for 24,500 iterations to track the 3×2 calibration grid to generate a pixel to millimeter conversion for every video recording. The training dataset was prepared by adding 5 videos with an average of 735 overall frames and extracting an average of 40 frames per video to manually label the six grid points of interest. The dataset was shuffled and split 95:5 for training and testing, respectively; and using a confidence threshold of 0.7, we observed average test error of 3.47 pixels and average train error of 3.42 pixels, compared to human provided annotations.

For real-time tracking of hindlimb body parts, a DeepLabCut model was trained using a MobileNet backbone. A training dataset was prepared by adding 6 videos recorded at 60, 125, and 415 FPS with an average of 887 overall frames and extracting an average of 100 frames per video to manually label the five anatomical landmarks of interest. The dataset was shuffled and split 95:5 for training and testing, respectively. The network was trained for 400,000 iterations using the training subset, and then evaluated on the held-out testing subset. Using a confidence threshold of 0.7, we observed average test error of 1.73 pixels and average train error of 1.59 pixels, compared to human provided annotations. Using the export model function within DLC, this model was exported into the Protocol Buffer format (.pb file) for seamless integration into Bonsai.

### 2.6 Offline pose analysis

Outputs from DLC were filtered and confidence thresholded. Pixel coordinates were converted to millimeters using calibration information obtained by a separate DLC model. The tracked body parts include the iliac crest (IC), hip, ankle, metatarsophalangeal joint (MTP), and the tip of the second toe. The position of the knee was inferred by measuring the lengths of the femur and tibia bones and then triangulation of the knee position using a custom matlab script.

Step cycle was analyzed using custom matlab scripts (https://github.com/tischfieldlab/Closed_Loop_Opto_Stimulation). In brief, local extrema positions of the toe were used to identify phase boundaries, with the local maxima of the toe indicating the start of the stance phase and the local minima of the toe indicating the start of the swing phase.

### 2.7 Hardware

To record and quantify hindlimb kinematics with real-time manipulation of neural activity during treadmill locomotion, six primary sets of hardware were utilized: a high performance Windows machine, openCV compatible high-speed camera, infrared lighting, motorized treadmill, pulse generator, and optogenetics supplies (**Table 1**). An openCV compatible camera is required to interface with Bonsai on a Windows machine. In this study, two cameras were purchased, one to record at high frame rates for offline pose estimation and a second with openCV compatibility for real-time pose estimation.

### 2.8 Software

All software tools have been compiled in **Table 2**. DeepLabCut-Live has three modes of operation, a stand alone GUI (DeepLabCut-Live! GUI), or pretrained model integration into Bonsai or Autopilot. In this study, we coupled a pre-trained DLC model with Bonsai to enable a closed-loop feedback system that detects a pose, performs an operation, and returns a processed pose (Kane et al., 2020).

**Table 2.** Software specifications for offline and real-time pose estimation as outlined in blue.

| Software | Function |
| --- | --- |
| **AOS Technologies AG Imaging Studio software suite** | Comprehensive camera suite to control camera capture settings and playback recorded videos |
| **Anaconda™** | Python distribution used to install DeepLabCut |
| **DeepLabCut™** | Open source pose estimation tool using for behavioral tracking |
| **Basler pylon camera software suite** | Comprehensive camera suite to control camera capture settings and playback recorded videos |
| **Bonsai** | Event based framework enabling acquisition and online processing of multiple data streams |
| **Matlab ®** | Programming language used for post processing and data analysis |

## 3. Results

### 3.1 Experimental setup and validation of pose tracking

Prior to data collection, animals were anesthetized, shaved on their right hindlimb area, and markers placed using white oil-based markers on five anatomical landmarks: iliac crest (IC), hip, ankle, metatarsophalangeal joint (MTP), and the tip of the second toe (**Figure 2A**). Animals were allowed to recover from anesthetic and then trained to locomote on the treadmill apparatus. Once trained to walk at speeds of 20 cm/s, we proceeded to capture high-speed video of the animals.

**Figure 2.**
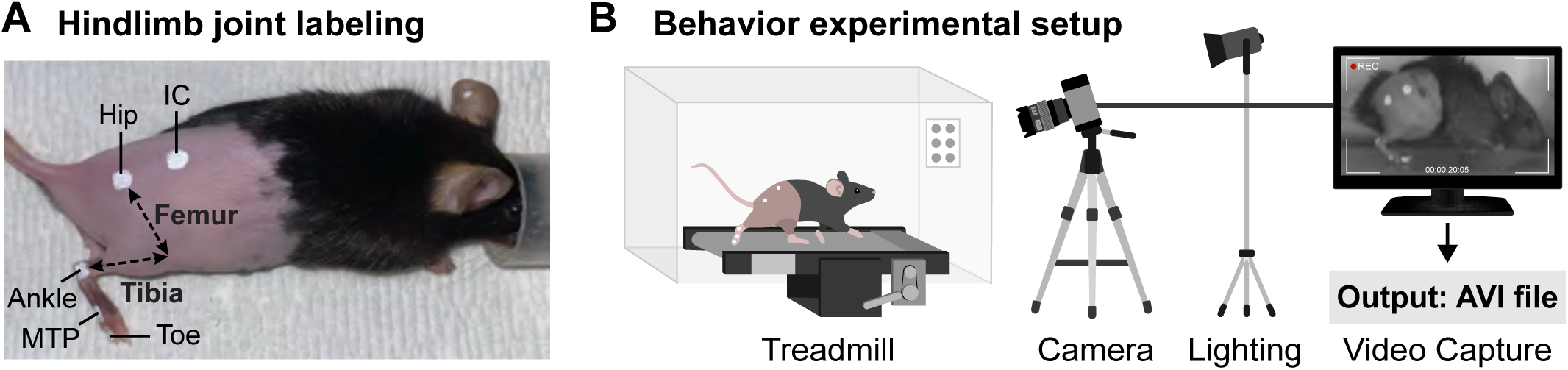
Experimental setup for video recording. **(A)** Prior to recording, labels are placed over five anatomical landmarks: iliac crest (IC), hip, ankle, metatarsophalangeal joint (MTP), and the tip of the second toe. The tibia and femur length are measured to triangulate the knee joint post-hoc. **(B)** A camera is mounted perpendicular to the treadmill to record sequences of images of an animal performing the task. In each recording, a calibration grid is placed in the field of view to convert pixels to metric units post-hoc. Recordings are stored on a computer as AVI files for post-hoc kinematic analysis.

To acquire video of a mouse walking on a treadmill, we used a motorized treadmill, and positioned a tripod mounted camera, oriented perpendicular to the direction of treadmill movement, approximately 1.5 meter distance from the treadmill. To increase illumination of the scene, while minimizing animal discomfort, we positioned two infrared spotlights on either side of the camera and oriented them to focus on the treadmill. In each video recording, a calibration grid was placed in the field of view to convert pixels to metric units. The output of this behavioral experiment is a dataset of video files used for subsequent analysis (**Figure 2B**).

We collected 5 videos consisting of a range between 5 and 27 step cycles per mouse, across 5 mice. Each video was trimmed to capture a minimum of 5 consecutive step cycles, forming a dataset of 9 videos. From this initial set of videos, we prepared a dataset totalling 1125 frames extracted from the captured videos and each frame labeled by an expert annotator. We then trained a DeepLabCut model for 300,000 iterations to accurately infer the positions of the five labeled body parts. Evaluation of the body part coordinates inferred by the model showed that the model predictions were accurate, having a mean test error of 2.38 pixels on held-out data compared to human annotations (**Figure 3A**).

**Figure 3.**
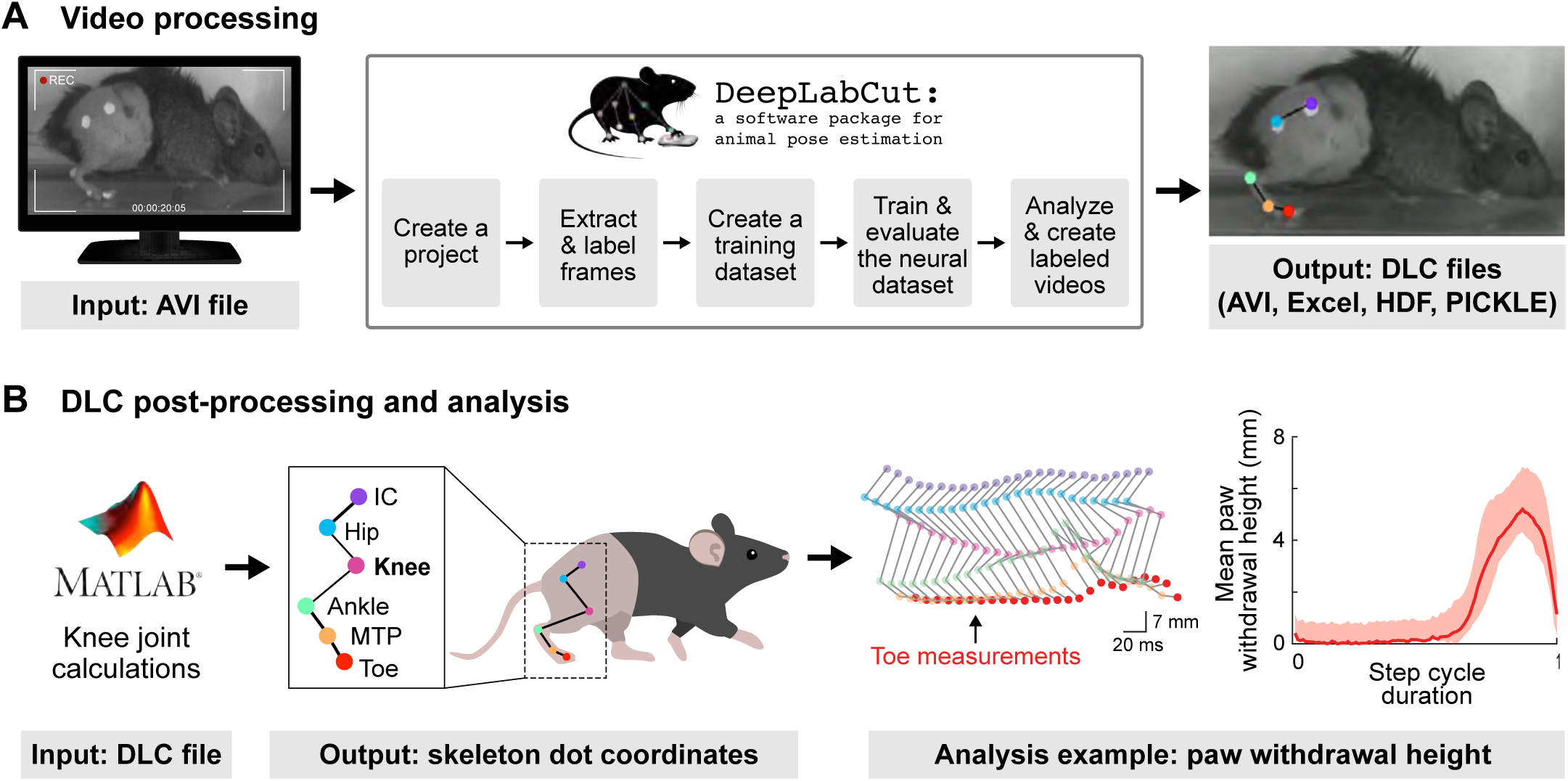
Pipeline for video analysis. **(A)** A DeepLabCut model is trained with manually labeled key points from a minimal subset of extracted video frames. Following training and testing, the DeepLabCut model produces and saves keypoint labels for every frame of the video recording for post-hoc analysis. **(B)** Using keypoint coordinates from (A), custom MATLAB scripts were generated to measure the X and Y coordinates of keypoints. Example data shows a stick plot with each keypoint labeled and the toe height tracked across a single step cycle (normalized 0-1).

### 3.2 Offline analysis and validation of step cycle and toe tracking

The ability to accurately detect step cycle boundaries as well as specific events within the step cycle is a critical requirement for this closed-loop intervention. This is also important for post-hoc analysis of experiments concerning the effect of such interventions. A given step cycle begins with the stance phase (the time between initial paw contact and liftoff), and progresses into the swing phase (the time between paw liftoff and contact with the surface again). To eliminate variability in step cycle durations that may confound comparisons within and between groups, step cycles were normalized from 0 to 1. We analyzed the step cycle in wild-type animals at a treadmill belt speed of 20 cm/s, with no fiber implant and observed an average step cycle duration to be 260 ms; dependent on the stride frequency of the individual mouse and consistent with previously published results (Leblond et al., 2003). We generated hindlimb skeletal stick plots to represent the progression of a step cycle using joint coordinates and phase boundaries. We also plotted the average trajectory of the toe coordinate over 10 consecutive normalized step cycles and found that in a normal animal, the toe height increased during the swing phase to an average peak height of 5.58 mm (SEM ± 1.25) (**Figure 3B**). This data provides a useful baseline characterization of the step cycle in wild-type animals without surgical manipulation.

### 3.3 Establish criteria for pose modulated stimulation

Offline analysis of step cycles from body part tracking data has the benefit of access to the entire time series of poses, however, a real-time closed-loop system only has access to the past and present (no future access). Additionally, offline analysis can tolerate heavier computational demands and long latency, while a real-time system must make fast decisions with only instantaneous pose information. We therefore set out to craft simple, low-latency, heuristics that accurately infer specific step cycle events from instantaneous pose data.

Using the complete hindlimb skeleton video recordings, we inspected videos within and between animals to craft rules for identifying specific phases of the step cycle, such as the initiation of stance, stance, the initiation of swing, and swing. In this study, differences in the x-coordinates of individual joints define phase criteria, given the assumption that the animal is always horizontally oriented with its nose pointing towards the right edge of the video and the tail pointing towards the left of the video and a coordinate system with the origin in the top right of the video. The initiation of the stance occurs when the X-position of the MTP is greater than the X-position of the IC. Stance occurs when the X-position of the ankle is greater than the X-position of the hip. The initiation of swing occurs when the X-position of the ankle is greater than the X-position of the MTP. Swing occurs when the X-position of the toe is greater than the X-position of the IC (**Figure 4A**).

**Figure 4.**
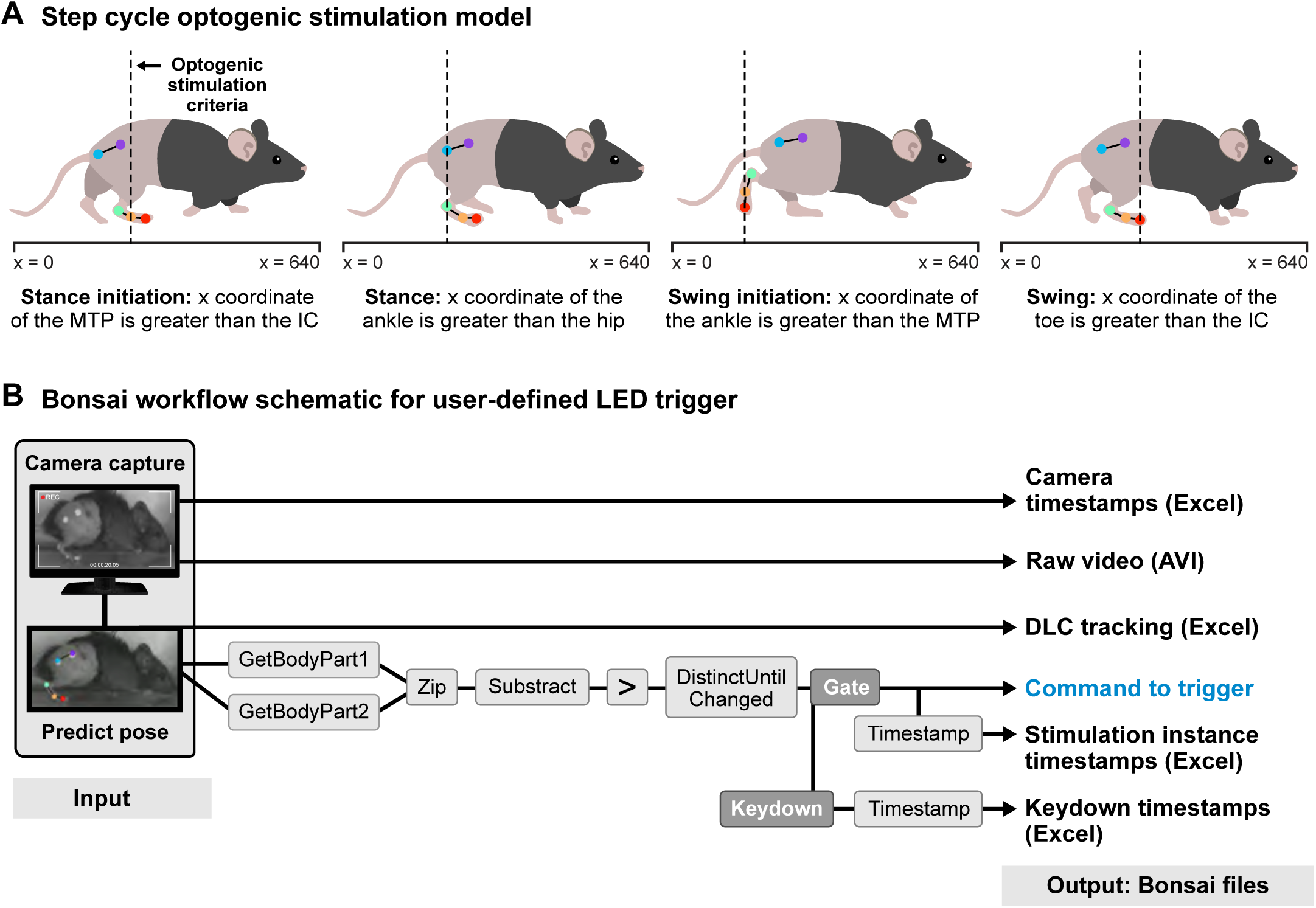
Pipeline for closed-loop system integration. **(A)** Video recordings are examined to establish user-defined thresholds for optogenetic stimulation. Here, the relative x-coordinates of keypoints were used to define specific phases of the step cycle, namely: Stance initiation, Stance, Swing initiation, and swing. **(B)** Bonsai workflow for user-defined optogenetic stimulation. Bonsai uses camera captured video acquisition coupled with a pre-trained DeepLabCut model to predict body part coordinates in real time. Using these coordinates, user-defined instances (A) are detected to trigger optogenetic stimulation.

### 3.4 Closed-loop system integration

We next modified our experimental apparatus to support a closed-loop stimulation experimental paradigm by adding hardware for optogenetic stimulation. First, a PulsePalv2 was connected to the acquisition computer via a USB cable. The digital output of the Pulse Pal was then connected to an LED driver, which was then connected to an LED. The output of the LED was routed via a fiber optic cable to the treadmill arena, where it terminated in a rotary joint. From there a fiber optic patch cable can be connected between the rotary joint and the cannula previously implanted above the spinal cord.

To capture and control the camera acquisition, we used the Basler Ace 2: a2A1920-160umPRO camera and the Basler pylon camera suite to preview camera settings prior to real-time pose estimation acquisition. To analyze incoming data and make the decision to stimulate or not, we programmed an experimental workflow in the Bonsai-RX environment with the following packages: DeepLabCut Library, DeepLabCut Design Library, Pylon Library, PulsePal Library, PulsePal Design Library. First, a *Camera Capture* node was configured to acquire images from the Basler camera at 150 FPS, and the images were routed to a *Video Writer* node to save the raw video stream to an AVI video file. Images from the *Camera Capture* node were also routed to a *PredictPose* node, which submits the input images to a DeepLabCut model for pose estimation and outputs the detected body part coordinates. Predicted body part coordinates were routed to a *CsvWriter* node, which writes these to a CSV file for later analysis. *GetBodyPart* nodes receive pose estimations and pick coordinates corresponding to the selected body parts (e.g. IC and MTP), selecting the X coordinate via *Position.X* nodes, which are combined via a *Zip* node and then subtracted from one another through a *Subtract* node. The result is converted to a boolean value through a *GreaterThan* node, comparing the result of the subtraction to zero, and then inverting the boolean result by comparison to False in an *Equal* node. To prevent multiple triggers (i.e. each frame after a condition is met until the condition is no longer met), the output is filtered by a *DistinctUntilChanged* node, which only produces distinct contiguous results. To allow the experimenter some control over valid experimental periods where stimulation may be allowed to be triggered, a *Gate* node, paired with a *KeyDown* node to allow triggering only within 30 seconds after the experimenter has pressed a keyboard key. When a sample is allowed through the gate, it is routed to a *TriggerOutput* node which triggers the Pulse Pal to begin playing the stimulation sequence, and the timestamp of the gate signal is recorded via a *CsvWriter* node (**Figure 4B**).

For stimulation targeted to different step cycle events (e.g. Stance initiation, stance, swing initiation, swing), we simply change the body parts compared by changing the body part name parameter in each of the two *GetBodyPart* nodes. Here, we collected a total of 16 videos, with 4 videos per mouse for each step cycle stimulation event, across 4 mice. Each video had a range between 10 - 14 single pulse stimulation triggers.

### 3.5 Validation of closed-loop LED triggering

Closed-loop systems have strict latency requirements in order to ensure desired interventions can be delivered in the proper moment. Latency can arise from several sources (e.g. Camera, pose estimation, pose analysis, stimulus delivery) and latency is additive across the experimental workflow. Additionally, the developed workflow must be effective and robust in detection of desired step cycle events and subsequent triggering of stimulus delivery.

To validate the effectiveness of the Bonsai workflow, a pilot study was designed to test the accuracy and latency of real-time DLC tracking and TriggerOutput commands once the optogenetic stimulation criteria is satisfied (**Figure 5A**). A treadmill-trained wild-type mouse was placed on the motorized treadmill and the fiber-coupled LED was taped to the side panel of the treadmill, in view of the camera (**Figure 5B**). Video recordings captured the mouse completing at least ten consecutive step cycles. Within the workflow, the *Gate* node recorded each instance that the gate opened to enable LED stimulation via *TriggerOuptut* once a key was pressed on the keyboard and the stimulation criteria was satisfied. Stimulation accuracy was then verified using the video file and excel sheet outputs of Bonsai to coordinate gate open timestamps with camera capture timestamps and aligning these timestamps to their corresponding frames in the video file (**Figure 5C**).

**Figure 5.**
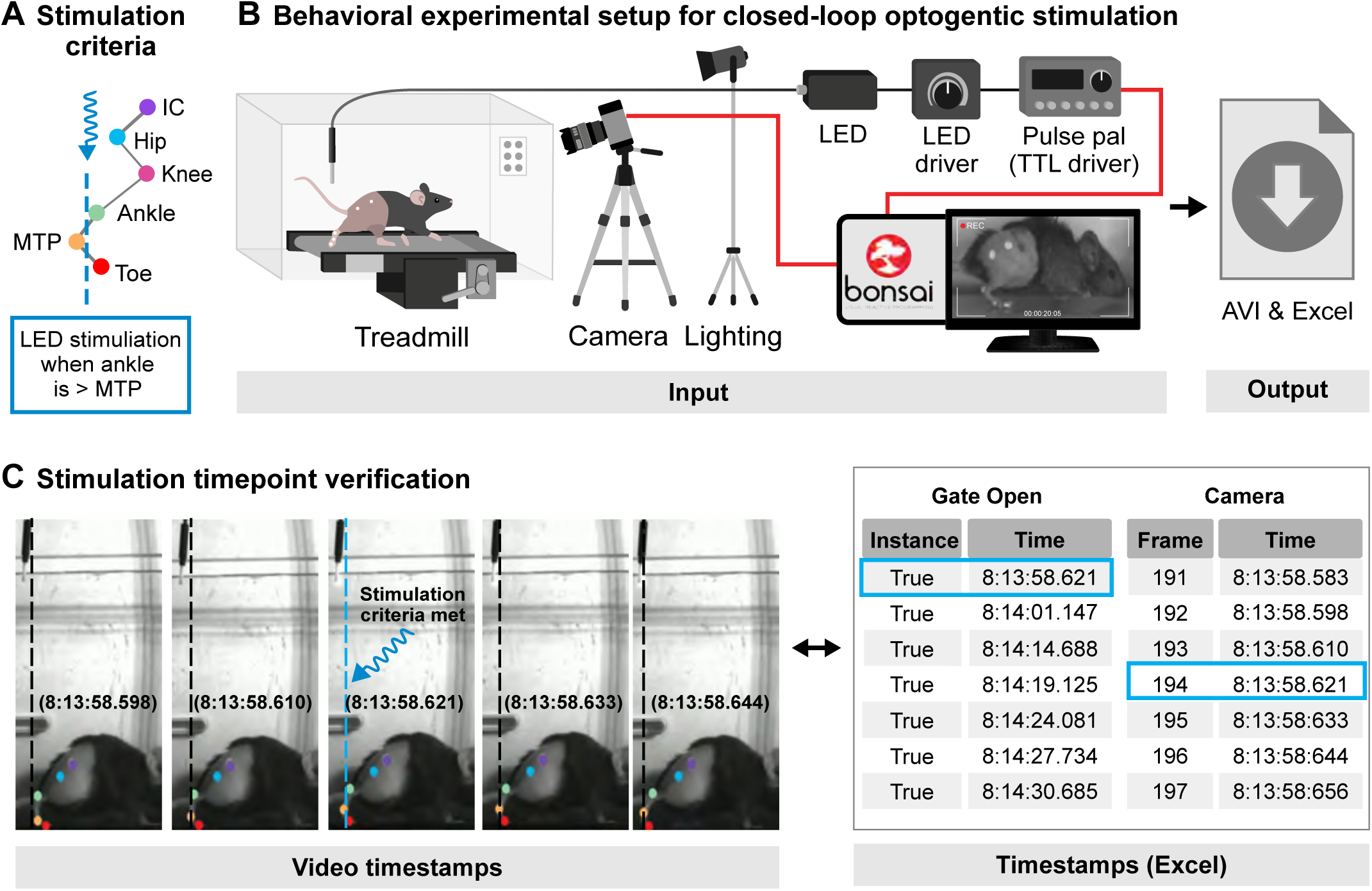
Validation of closed-loop triggering. **(A)** Example user-defined instance to identify swing initiation (Ankle > MTP). **(B)** Behavioral setup is as in Figure 1, with the addition of an optic fiber in the camera field of view. Here, the bonsai workflow is integrated to the behavioral setup to provide an external command to a TTL driver that triggers LED activation at user-defined instances. **(C)** Example visual inspection of closed-loop optogenetic stimulation at a swing initiation. Right: Instance detection of the open Gate and camera timestamps are then compared to validate closed-loop triggering.

Visual inspection to align when the LED is ON with an instance such as the initiation of swing when the x-coordinate of the ankle exceeds the MTP yields a latency dependent on the frame rate. At 150 FPS, a frame is generated every 6.67 ms and the LED may turn ON within this time interval before it is visible in the video frame. True latency is determined by calculating the latency of real-time DLC tracking, overall system latency such as computer processing speeds and Pulse Pal trigger output speeds, and subtracting the delay between the instance detection and the command to trigger the LED. This first pass testing corroborates that the overall system is effective at triggering an LED to turn ON at pose-modulated instances and indicates its potential use to photo stimulate genetically encoded tools in mice.

### 3.6 Validation of optogenetic activation of TRPV1^+^ primary afferent fibers

To validate our ability to successfully activate nociceptive primary afferent fibers, we anesthetized TRPV1^Cre^;Advillin^FlpO^;R26^LSL-FSF-ChR2^ ^(Ai80)^ mice and applied direct photostimulation (2.2 mW, 10 ms pulses, 10 Hz for 30 min) to the spinal cord by positioning an optic fiber probe (400 nm core, 1 mm fiber length, ThorLabs) above the spinal cord surface. Animals were maintained under anesthesia, then perfused transcardially with 4% paraformaldehyde 1hr following photostimulation. Spinal cord sections from photostimulation segments were processed and immunolabeled to visualize EYFP (to label TRPV1-ChR2^+^ afferent terminals), the activity marker c-Fos (to label activated spinal cord neurons), and NeuN (to label dorsal horn neurons). In line with previous characterizations of TRPV1^+^ sensory afferents (Caterina et al., 1999, 2000; Samineni et al., 2017), EYFP expression was largely restricted to the superficial dorsal horn (LI-LII, **Figure 1B**). As expected from the location of EYFP^+^ terminals, we observed a significant increase in c-Fos^+^ neurons within LI-II ipsilateral to photostimulation (**Figure 1B-C**). Together, this data demonstrates our ability to optogenetically stimulate TRPV1^+^ primary afferent fibers within the dorsal horn of the spinal cord.

### 3.7 Characterizing the effects from photostimulation during specific step cycle events

We finally sought to evaluate the effects of *in vivo* photostimulation of TRPV1^+^ primary afferent fibers in the dorsal horn of the spinal cord in awake, behaving mice at different phases of the step cycle. TRPV1^+^ primary afferent fibers are known to transmit nociceptive information, and their activation has previously been shown to evoke nociceptive responses (i.e. withdrawal of the paw in response to pain) (Beaudry et al., 2017; Samineni et al., 2017). We hypothesized that stimulation of these neurons would cause reflexive paw withdrawal, and interrupt the normal progression of step cycle dynamics.

To test this hypothesis, TRPV1^Cre^;Advillin^FlpO^;R26^LSL-FSF-ChR2(Ai80)^ animals were surgically implanted with a fiber optic probe positioned to illuminate the dorsal horn of the spinal cord at the L3-4 level (**Figure 1A**). Mice were allowed to recover and subsequently trained to walk on the treadmill at a belt speed of 20 cm/s. On the day of experimentation, mice were habituated to the behavior room, and gently placed on the treadmill. Using the Bonsai workflow described in 3.4, we recorded animals walking while simultaneously evaluating the poses of the right hindlimb. After an animal successfully performed at least 5 step cycles, the experimenter pressed a *KeyDown* to allow the *Gate* node to enable pose-triggered photostimulation at the chosen step cycle event. Photostimulation consisted of a single 2.2 mW 10 ms pulse. The data collected was then processed offline for analysis of step cycle and paw withdrawal responses.

As expected, animals were responsive to the photoactivation of TRPV1^+^ sensory afferents during treadmill locomotion (**Figure 6A**). Stick plots visualize the trajectory of each hindlimb joint during each stimulation event and highlight the elevation of the paw following stimulation and during the swing phase (**Figure 6B**). Variability between stick plot representations are attributed to a subject’s individual stepping frequency and differences in responses between the step cycle stimulation events (i.e. prolonged paw elevation as opposed to brief responses).

**Figure 6.**
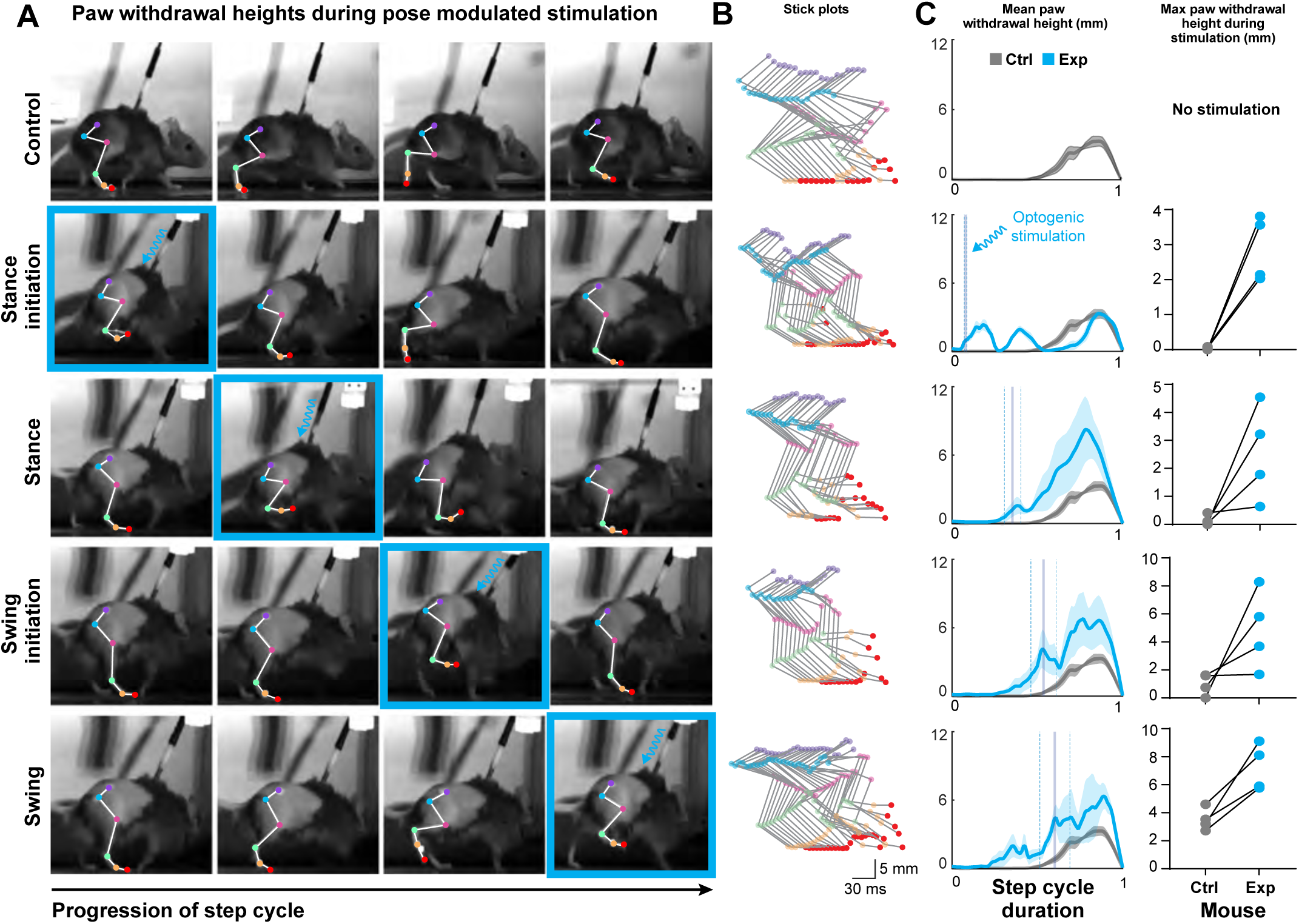
Closed-loop optogenetic stimulation of nociceptive primary afferents during locomotion. **(A)** TRPV1^Cre^;Advillin^FlpO^;^R26LSL-FSF-ChR2(Ai80)^ mice were implanted with an optic probe at the L3-4 spinal cord level. During treadmill locomotion (20 cm/s) user-defined instances were used to trigger optogenetic stimulation at specific phases of the step cycle: Control – no stimulation; Stance initiation; Stance; Swing initiation; Swing. Timing of optogenetic stimulation is depicted with a blue outline surrounding the frame. **(B)** Representative stick plots for each instance with key points labeled. **(C)** Quantification of paw height tracked across the step cycle (normalized 0-1) and maximum paw withdrawal height in control (gray, no optogenetic stimulation) and experimental (blue, optogenetic stimulation) steps. The average optogenetic stimulation time and SEM are indicated by vertical lines. Paired T-tests; Stance initiation, **p = 0.0097; Stance, ns p = 0.0804; Swing initiation, ns p = 0.1192; Swing, *p = 0.0170. n = 4 mice.

Peak paw withdrawal is defined as the highest paw elevation response following photo stimulation within the defined step cycle event. With a treadmill belt speed of 20 cm/s, by the time the right hindlimb strikes the ground to initiate stance, the left hindlimb is grounded and preparing to initiate swing. During optogenetic stimulation at the initiation of stance, the mice generated a brief paw withdrawal with an average elevation peak of 2.89 mm (SEM ± 0.46), compared to the average 0.03 mm (SEM ± 0.02) without stimulation. The subjects consistently responded with an elevation of the right hindlimb, and on many occasions, the mice used their left hindlimb to propel themselves forward, as if to normally start the swing phase, and then both hind paws were briefly elevated in the air. This reaction frequently caused the mice to generate a second paw elevation, as seen with the first two peaks in paw withdrawal heights, prior to readjusting their step and continuing with a more normal swing phase. During stimulation of the stance phase, the mice generated an average peak paw withdrawal height of 2.54 mm (SEM ± 0.85), compared to the average 0.14 mm (SEM ± 0.09) without stimulation. This stimulation generated variable responses in which the mice may briefly elevate their right paw and the paw stutters before coming in contact with the ground, or the right paw exhibits a large withdrawal elevation and once it comes in contact with the ground, the mouse is prepared to continue with normal step cycles (**Figure 6B-C**).

At 20 cm/s, when the right hindlimb is preparing to initiate swing, the left hindlimb is grounded because it recently initiated stance. Optogenetic stimulation at the initiation of swing generates an initial average paw withdrawal peak height of 4.86 mm (SEM ± 1.42), compared to the average 0.99 mm (SEM ± 0.39) without stimulation. During this stimulation event, the mice frequently responded with increased knee flexion and an elevated right hindlimb, followed by an elevation of the left hindlimb, so that both hind paws are in the air until either hindlimb contacts the ground (**Figure 6B-C**). The combination of the right and left hindlimb elevation increased the overall response time. Photo stimulation within the stance phase and at the initiation of the swing phase generate similar responses with elevation during the stimulation period, followed by an extended duration and more elevated paw position prior to transitioning into a normal step. Lastly, during stimulation of the swing phase, the mice generated an average peak paw withdrawal elevation of 7.22 mm (SEM ± 0.83), compared to the average 3.51 mm (SEM ± 0.39) without stimulation. The first hump of the withdrawal elevation is the onset of a normal swing phase and the following peaks are the elevations that occur due to stimulation (**Figure 6C**). Variation in the stimulation timing may be attributed to differences in stride frequency. Overall, these responses highlight the phase dependence of nociceptive withdrawal responses, and though not quantified here, point to an important role for left-right hindlimb coordination.

## 4. Discussion

### 4.1 A Novel Closed-Loop System for Probing State-Dependent Neural Circuits

Our study demonstrates a novel closed-loop system that integrates real-time pose estimation with optogenetic manipulation to probe state-dependent neural circuits during locomotion. By combining DeepLabCut-Live pose tracking with phase-specific optogenetic stimulation of nociceptive sensory neurons, we were able to begin to investigate how sensory inputs modulate locomotor output in a context-dependent manner. This approach addresses a key challenge in studying adaptive locomotor behaviors by allowing precise temporal control of neural manipulation based on the ongoing motor state.

### 4.2 Phase-Dependent Effects of Sensory Stimulation on Locomotor Output

The results from our proof-of-principle experiments with TRPV1^+^ sensory afferent stimulation reveal intriguing phase-dependent effects on locomotor output. Photostimulation during both stance and swing phases initiated a paw withdrawal response, but the characteristics of these responses differed markedly depending on the phase of stimulation. When stimulation occurred during stance, with the paw on the ground, it elicited a brief withdrawal response with an average peak height of 2.54 mm (**Figure 6C**). In contrast, stimulation during swing, when the paw was already in the air, produced a more pronounced response. This swing-phase stimulation led to an extended withdrawal with an average peak height of 7.22 mm and was accompanied by increased knee flexion (**Figure 6B-C**).

These phase-dependent differences in response magnitude and kinematics align with our understanding of how sensory input is processed differently depending on the current state of the limb. The more pronounced response during swing, characterized by greater elevation and increased knee flexion, may reflect the engagement of protective reflexes similar to the stumbling corrective response (Mayer & Akay, 2018). This observation underscores the importance of state-dependent sensorimotor integration in shaping adaptive locomotor behaviors.

### 4.3 Bilateral Responses to Unilateral Stimulation: Implications for Spinal Circuit Coordination

Intriguingly, while our optogenetic stimulation was unilateral, as confirmed by c-Fos immunostaining (**Figure 1B**), we observed bilateral responses in the hindlimbs during stance and swing stimulations. Though we only tracked and quantified right hindlimb kinematics, visual inspection of video recordings demonstrated an initial withdrawal occurring in the stimulated (right) hindpaw, followed by elevation of the contralateral (left) hindpaw (**Figure 6A**).This bilateral response to unilateral stimulation highlights the intricate left-right coordination mechanisms within the spinal locomotor circuitry and suggests that sensory inputs can modulate locomotor patterns across both sides of the spinal cord, likely in a stimulation intensity and phase-dependent manner.

Indeed, previous work has shown that motor responses of one limb can be initiated by cutaneous stimulation applied to the contralateral limb (Gauthier & Rossignol, 1981; Perl, 1957). These contralateral responses are mediated by spinal cord commissural neurons that are responsive to cutaneous stimulation (Laflamme et al., 2023), as well as descending serotonergic modulation (Abbinanti et al., 2012; Butt & Kiehn, 2003). The phase-dependent nature of this bilateral response further underscores the context-specific integration of sensory information within the locomotor circuitry, which may be influenced by both local spinal circuits and descending control.

### 4.4 Future Applications and Potential Enhancements of the System

The ability to manipulate specific neuronal populations at precise phases of the step cycle opens new avenues for investigating the neural control of locomotion. Our system’s capacity to elicit and measure these phase-dependent and bilaterally coordinated responses demonstrates its utility in probing the complex interactions within spinal circuits during ongoing behavior. Future studies could leverage this approach to further dissect the neural mechanisms underlying interlimb coordination, potentially by combining our technique with physiological recordings of muscle (Chung et al., 2023; Pearson et al., 2005) and/or spinal interneurons (Lavaud et al., 2024). Moreover, this system’s versatility allows for broader applications in studying state-dependent behaviors. The real-time pose estimation component also offers the potential to trigger manipulations based on more complex behavioral events or postures, extending beyond simple phase-based criteria.

To further enhance the system’s capabilities and reduce latency, several improvements could be implemented. Incorporating a forward prediction filter, such as a Kalman filter, could help compensate for processing delays by anticipating future poses (Kane et al., 2020). Optimizing camera frame rates and resolution, as well as utilizing high-performance computing hardware, can minimize overall system latency. Additionally, expanding to multi-camera setups to acquire 3D motion analysis would provide a more comprehensive view of the animal’s behavior and potentially improve the accuracy of pose estimation and phase detection by mitigating occlusions.

### 4.5 Conclusion: Bridging Cellular Manipulations with Complex Behaviors

Our integrated DeepLabCut-Live (Kane et al., 2020) and optogenetics approach provides a powerful new tool for probing the neural circuits underlying adaptive motor behaviors. By enabling precise, state-dependent manipulation of neural activity, this system helps bridge the gap between cellular-level manipulations and complex behavioral outputs. The phase-dependent and bilaterally coordinated responses we observed highlight the complex nature of sensorimotor integration during locomotion and demonstrate the potential of this approach in unraveling the intricacies of motor control. As the field of neuroscience continues to emphasize the importance of studying neural circuits in the context of naturalistic behaviors, tools like the one presented here will be crucial in advancing our understanding of how the nervous system generates and modulates adaptive behaviors.

## Acknowledgements

Financial support was provided by Whitehall Foundation (to V.E.A.), Craig H. Neilsen Foundation (to V.E.A.), Pew Charitable Trust (to V.E.A.), NJ Commission on Spinal Cord Research (to M.A.G., and V.E.A.), NIH NINDS K01NS116224 (to V.E.A.), NIH NINDS R01NS119268 (to V.E.A.), NIH NINDS R01NS124799 (to V.E.A) and NIH NINDS R01NS119268-S1 (to M. Gonzalez).

## Author contributions

Behavior experiments, M.Gonzalez and M.A.Gradwell; Behavior analysis, M.Gonzalez with help from M.A.Gradwell and J.K.Thackray; Anatomy experiments, M.A.Gradwell with help from K.R.Patel and K.K.Temkar; Preparation of figures, M.Gonzalez and M.A.Gradwell with help from simplifiedsciencepublishing; Experiment design, M.Gonzalez and M.A.Gradwell; Study conception, V.E.Abraira; Writing, M.Gonzalez with help from M.A.Gradwell, J.K.Thackray, and V.E.Abraira.

## Declaration of generative AI and AI-assisted technologies in the writing process

During the preparation of this work the author(s) used Claude3 in order to improve the clarity and readability of the manuscript. After using this tool/service, the author(s) reviewed and edited the content as needed and take(s) full responsibility for the content of the publication.

## References

Abbinanti, M. D., Zhong, G., & Harris-Warrick, R. M. (2012). Postnatal emergence of serotonin-induced plateau potentials in commissural interneurons of the mouse spinal cord. Journal of Neurophysiology, 108(8), 2191–2202.

Beaudry, H., Daou, I., Ase, A. R., Ribeiro-da-Silva, A., & Séguéla, P. (2017). Distinct behavioral responses evoked by selective optogenetic stimulation of the major TRPV1+ and MrgD+ subsets of C-fibers. Pain, 158(12), 2329–2339.

Boyden, E. S., Zhang, F., Bamberg, E., Nagel, G., & Deisseroth, K. (2005). Millisecond-timescale, genetically targeted optical control of neural activity. Nature Neuroscience, 8(9), 1263–1268.

Butt, S. J. B., & Kiehn, O. (2003). Functional identification of interneurons responsible for left-right coordination of hindlimbs in mammals. Neuron, 38(6), 953–963.

Caterina, M. J., Leffler, A., Malmberg, A. B., Martin, W. J., Trafton, J., Petersen-Zeitz, K. R., Koltzenburg, M., Basbaum, A. I., & Julius, D. (2000). Impaired nociception and pain sensation in mice lacking the capsaicin receptor. Science, 288(5464), 306–313.

Caterina, M. J., Rosen, T. A., Tominaga, M., Brake, A. J., & Julius, D. (1999). A capsaicin-receptor homologue with a high threshold for noxious heat. Nature, 398(6726), 436–441.

Cavanaugh, D. J., Chesler, A. T., Jackson, A. C., Sigal, Y. M., Yamanaka, H., Grant, R., O’Donnell, D., Nicoll, R. A., Shah, N. M., Julius, D., & Basbaum, A. I. (2011). Trpv1 reporter mice reveal highly restricted brain distribution and functional expression in arteriolar smooth muscle cells. The Journal of Neuroscience: The Official Journal of the Society for Neuroscience, 31(13), 5067–5077.

Chung, B., Zia, M., Thomas, K. A., Michaels, J. A., Jacob, A., Pack, A., Williams, M. J., Nagapudi, K., Teng, L. H., Arrambide, E., Ouellette, L., Oey, N., Gibbs, R., Anschutz, P., Lu, J., Wu, Y., Kashefi, M., Oya, T., Kersten, R.,…Sober, S. J. (2023). Myomatrix arrays for high-definition muscle recording. eLife, 12. 10.7554/eLife.88551

Chuong, A. S., Miri, M. L., Busskamp, V., Matthews, G. A. C., Acker, L. C., Sørensen, A. T., Young, A., Klapoetke, N. C., Henninger, M. A., Kodandaramaiah, S. B., Ogawa, M., Ramanlal, S. B., Bandler, R. C., Allen, B. D., Forest, C. R., Chow, B. Y., Han, X., Lin, Y., Tye, K. M.,…Boyden, E. S. (2014). Noninvasive optical inhibition with a red-shifted microbial rhodopsin. Nature Neuroscience, 17(8), 1123–1129.

Daigle, T. L., Madisen, L., Hage, T. A., Valley, M. T., Knoblich, U., Larsen, R. S., Takeno, M. M., Huang, L., Gu, H., Larsen, R., Mills, M., Bosma-Moody, A., Siverts, L. A., Walker, M., Graybuck, L. T., Yao, Z., Fong, O., Nguyen, T. N., Garren, E.,…Zeng, H. (2018). A Suite of Transgenic Driver and Reporter Mouse Lines with Enhanced Brain-Cell-Type Targeting and Functionality. Cell, 174(2), 465–480.e22.

Deisseroth, K. (2011). Optogenetics. Nature Methods, 8(1), 26–29.

Engberg, I., & Lundberg, A. (1969). An electromyographic analysis of muscular activity in the hindlimb of the cat during unrestrained locomotion. Acta Physiologica Scandinavica, 75(4), 614–630.

Forssberg, H. (1979). Stumbling corrective reaction: a phase-dependent compensatory reaction during locomotion. Journal of Neurophysiology, 42(4), 936–953.

Forssberg, H., Grillner, S., & Rossignol, S. (1977). Phasic gain control of reflexes from the dorsum of the paw during spinal locomotion. Brain Research, 132(1), 121–139.

Gauthier, L., & Rossignol, S. (1981). Contralateral hindlimb responses to cutaneous stimulation during locomotion in high decerebrate cats. Brain Research, 207(2), 303–320.

Grillner, S., & El Manira, A. (2020). Current Principles of Motor Control, with Special Reference to Vertebrate Locomotion. Physiological Reviews, 100(1), 271–320.

Harnie, J., Audet, J., Mari, S., Lecomte, C. G., Merlet, A. N., Genois, G., Rybak, I. A., Prilutsky, B. I., & Frigon, A. (2022). State-and Condition-Dependent Modulation of the Hindlimb Locomotor Pattern in Intact and Spinal Cats Across Speeds. Frontiers in Systems Neuroscience, 16, 814028.

Hughes, D. I., Sikander, S., Kinnon, C. M., Boyle, K. A., Watanabe, M., Callister, R. J., & Graham, B. A. (2012). Morphological, neurochemical and electrophysiological features of parvalbumin-expressing cells: a likely source of axo-axonic inputs in the mouse spinal dorsal horn. The Journal of Physiology, 590(16), 3927–3951.

Kane, G. A., Lopes, G., Saunders, J. L., Mathis, A., & Mathis, M. W. (2020). Real-time, low-latency closed-loop feedback using markerless posture tracking. eLife, 9. 10.7554/eLife.61909

Kiehn, O. (2016). Decoding the organization of spinal circuits that control locomotion. Nature Reviews. Neuroscience, 17(4), 224–238.

Kiehn, O., Dougherty, K. J., Hägglund, M., Borgius, L., Talpalar, A., & Restrepo, C. E. (2010). Probing spinal circuits controlling walking in mammals. Biochemical and Biophysical Research Communications, 396(1), 11–18.

Klapoetke, N. C., Murata, Y., Kim, S. S., Pulver, S. R., Birdsey-Benson, A., Cho, Y. K., Morimoto, T. K., Chuong, A. S., Carpenter, E. J., Tian, Z., Wang, J., Xie, Y., Yan, Z., Zhang, Y., Chow, B. Y., Surek, B., Melkonian, M., Jayaraman, V., Constantine-Paton, M.,…Boyden, E. S. (2014). Independent optical excitation of distinct neural populations. Nature Methods, 11(3), 338–346.

Laflamme, O. D., Markin, S. N., Deska-Gauthier, D., Banks, R., Zhang, Y., Danner, S. M., & Akay, T. (2023). Distinct roles of spinal commissural interneurons in transmission of contralateral sensory information. Current Biology: CB, 33(16), 3452–3464.e4.

Lavaud, S., Bichara, C., D’Andola, M., Yeh, S.-H., & Takeoka, A. (2024). Two inhibitory neuronal classes govern acquisition and recall of spinal sensorimotor adaptation. Science, 384(6692), 194–201.

Leblond, H., L’Esperance, M., Orsal, D., & Rossignol, S. (2003). Treadmill locomotion in the intact and spinal mouse. The Journal of Neuroscience: The Official Journal of the Society for Neuroscience, 23(36), 11411–11419.

Lopes, G., Bonacchi, N., Frazão, J., Neto, J. P., Atallah, B. V., Soares, S., Moreira, L., Matias, S., Itskov, P. M., Correia, P. A., Medina, R. E., Calcaterra, L., Dreosti, E., Paton, J. J., & Kampff, A. R. (2015). Bonsai: an event-based framework for processing and controlling data streams. Frontiers in Neuroinformatics, 9, 7.

Mardinly, A. R., Oldenburg, I. A., Pégard, N. C., Sridharan, S., Lyall, E. H., Chesnov, K., Brohawn, S. G., Waller, L., & Adesnik, H. (2018). Precise multimodal optical control of neural ensemble activity. Nature Neuroscience, 21(6), 881–893.

Mathis, A., Mamidanna, P., Cury, K. M., Abe, T., Murthy, V. N., Mathis, M. W., & Bethge, M. (2018). DeepLabCut: markerless pose estimation of user-defined body parts with deep learning. Nature Neuroscience, 21(9), 1281–1289.

Mayer, W. P., & Akay, T. (2018). Stumbling corrective reaction elicited by mechanical and electrical stimulation of the saphenous nerve in walking mice. The Journal of Experimental Biology, 221(Pt 13). 10.1242/jeb.178095

Nath, T., Mathis, A., Chen, A. C., Patel, A., Bethge, M., & Mathis, M. W. (2019). Using DeepLabCut for 3D markerless pose estimation across species and behaviors. Nature Protocols, 14(7), 2152–2176.

Pearson, K. G., Acharya, H., & Fouad, K. (2005). A new electrode configuration for recording electromyographic activity in behaving mice. Journal of Neuroscience Methods, 148(1), 36–42.

Pereira, T. D., Tabris, N., Matsliah, A., Turner, D. M., Li, J., Ravindranath, S., Papadoyannis, E. S., Normand, E., Deutsch, D. S., Wang, Z. Y., McKenzie-Smith, G. C., Mitelut, C. C., Castro, M. D., D’Uva, J., Kislin, M., Sanes, D. H., Kocher, S. D., Wang, S. S.-H., Falkner, A. L.,…Murthy, M. (2022). SLEAP: A deep learning system for multi-animal pose tracking. Nature Methods, 19(4), 486–495.

Perl, E. R. (1957). Crossed reflexes of cutaneous origin. The American Journal of Physiology, 188(3), 609–615.

Rossignol, S. (1996). Neural control of stereotypic limb movements. In S. J. Rowell L (Ed.), Handbook of Physiology section 12 Exercise: Regulation and Integration of Multiple Systems. (pp. 173–216). American Physiological Society.

Roth, B. L. (2016). DREADDs for Neuroscientists. Neuron, 89(4), 683–694.

Samineni, V. K., Yoon, J., Crawford, K. E., Jeong, Y. R., McKenzie, K. C., Shin, G., Xie, Z., Sundaram, S. S., Li, Y., Yang, M. Y., Kim, J., Wu, D., Xue, Y., Feng, X., Huang, Y., Mickle, A. D., Banks, A., Ha, J. S., Golden, J. P.,…Gereau, R. W., 4th. (2017). Fully implantable, battery-free wireless optoelectronic devices for spinal optogenetics. Pain, 158(11), 2108–2116.

Sjulson, L., Cassataro, D., DasGupta, S., & Miesenböck, G. (2016). Cell-Specific Targeting of Genetically Encoded Tools for Neuroscience. Annual Review of Genetics, 50, 571–594.

Smith, K. M., Browne, T. J., Davis, O. C., Coyle, A., Boyle, K. A., Watanabe, M., Dickinson, S. A., Iredale, J. A., Gradwell, M. A., Jobling, P., Callister, R. J., Dayas, C. V., Hughes, D. I., & Graham, B. A. (2019). Calretinin positive neurons form an excitatory amplifier network in the spinal cord dorsal horn. eLife, 8. 10.7554/eLife.49190

Wand, P., Prochazka, A., & Sontag, K. H. (1980). Neuromuscular responses to gait perturbations in freely moving cats. Experimental Brain Research. Experimentelle Hirnforschung. Experimentation Cerebrale, 38(1), 109–114.

Yizhar, O., Fenno, L. E., Davidson, T. J., Mogri, M., & Deisseroth, K. (2011). Optogenetics in neural systems. Neuron, 71(1), 9–34.

Zimmerman, A. L., Kovatsis, E. M., Pozsgai, R. Y., Tasnim, A., Zhang, Q., & Ginty, D. D. (2019). Distinct Modes of Presynaptic Inhibition of Cutaneous Afferents and Their Functions in Behavior. Neuron, 102(2), 420–434.e8.

